# A distinct mechanism of RNA recognition by the transcription factor GATA1

**DOI:** 10.1101/2024.12.02.626266

**Authors:** Daniella A. Ugay, Robert T. Batey, Deborah S. Wuttke

## Abstract

Several human transcription factors (TFs) have been reported to directly bind RNA through non-canonical RNA-binding domains; however, most of these TFs remain to be further validated as *bona fide* RNA-binding proteins (RBPs). Our systematic analysis of RBP discovery datasets reveals a varied set of candidate TF-RBPs that encompass most TF families. These candidate RBPs include members of the GATA family, which are essential factors in embryonic development. Investigation of the RNA-binding features of GATA1, a major hematopoietic TF, reveals robust non-sequence specific binding to RNAs *in vitro*. Moreover, RNA binding by GATA1 is competitive with DNA binding, which occurs through a shared binding surface spanning the DNA-binding domain and arginine-rich motif (ARM) like domain. We show that the ARM-like domain contributes both substantially to high-affinity DNA binding and electrostatically to plastic RNA recognition, suggesting that the separable RBD assigned to the ARM-domain in GATA1 is an oversimplification of a more complex recognition network. These biochemical data demonstrate a unified integration of DNA- and RNA-binding surfaces within GATA1, whereby the ARM-like domain provides an electrostatic surface for RNA binding but does not fully dominate GATA1-RNA interactions, which may also apply to other TF-RBPs. This competitive DNA/RNA binding activity using overlapping nucleic acid binding regions points to the possibility of RNA-mediated regulation of GATA1 function during hematopoiesis. Our study highlights the multifunctionality of DNA-binding domains in RNA recognition and supports the need for robust characterization of predicted non-canonical RNA-binding domains such as ARM-like domains.

## Introduction

A growing number of high-throughput RNA-binding protein (RBP) discovery methods have expanded the repertoire of potential RBPs.^1^ These include human transcription factors (TFs), which have been shown to bind RNA via zinc finger (ZF) motifs and, more recently, through non-canonical RNA binding motifs such as an arginine-rich motif (ARM) like domain located adjacent to DNA-binding domains (DBDs).^2,3^ A prevailing model suggests that TFs interact with DNA and RNA through separate binding domains, whereby RNA binding via an ARM-like domain could enhance TF’s chromatin occupancy and modulate transcription.^3,4^

Though likely true for a set of RNA-binding TFs, the proposed model for simultaneous TF-DNA-RNA binding via separate domains may not be universal. We have previously shown that TFs from two major families, the nuclear hormone receptors glucocorticoid receptor (GR), estrogen receptor alpha (ERα), and high-mobility group box Sox2 bind DNA and RNA competitively, ruling out the possibility of functionally distinct surfaces for binding DNA and RNA.^5–9^ We further demonstrated that RNA can compete GR off chromatin, limiting dexamethasone-stimulated gene activation.^10^ The prevalence of this competitive DNA/RNA binding in other TF families and its functional consequences on transcription requires further investigation. Critically, newly discovered TF-RBPs require a rigorous experimental evaluation to validate TF-RNA interactions, define molecular characteristics of DNA/RNA binding, and establish the functional role of these interactions in transcriptional regulation.

## Results

### Identification of RNA-binding TFs spanning most TF families and selection of GATA1

To investigate the scope of DNA/RNA interacting TFs in the human proteome, we probed the RNA-binding potential in over 1600 human TFs by analyzing RBP annotations reported in twenty independent RBP discovery studies.^11–31^ Briefly, we combined these RBP datasets and matched RBP gene identifiers with those of cataloged human TFs.^11^ Overlapping hits were sorted for TFs with at least one RBP annotation, yielding 285 human TFs with a potential to associate with RNA (Figure 1A, Table S1), consistent with a recently published survey.^3^ We categorized these TF-RBPs into their respective DBD families and quantified the RBP annotations in each family. This analysis identified TF-RNA interactions in at least 32 DBD families (Figure 1B), some of which have demonstrated RNA binding activities (e.g., C2H2 ZFs),^32^ and others have not been systematically assessed yet (GATA, MBD, THAP finger, etc.). Thus, our analysis corroborates experimental evidence for TF-RNA binding^3^ and expands the TF-RBP repertoire, now spanning diverse DBDs and potentially involving unexplored mechanisms of RNA recognition beyond the canonical RNA-binding domains.^33^

**Figure 1.**
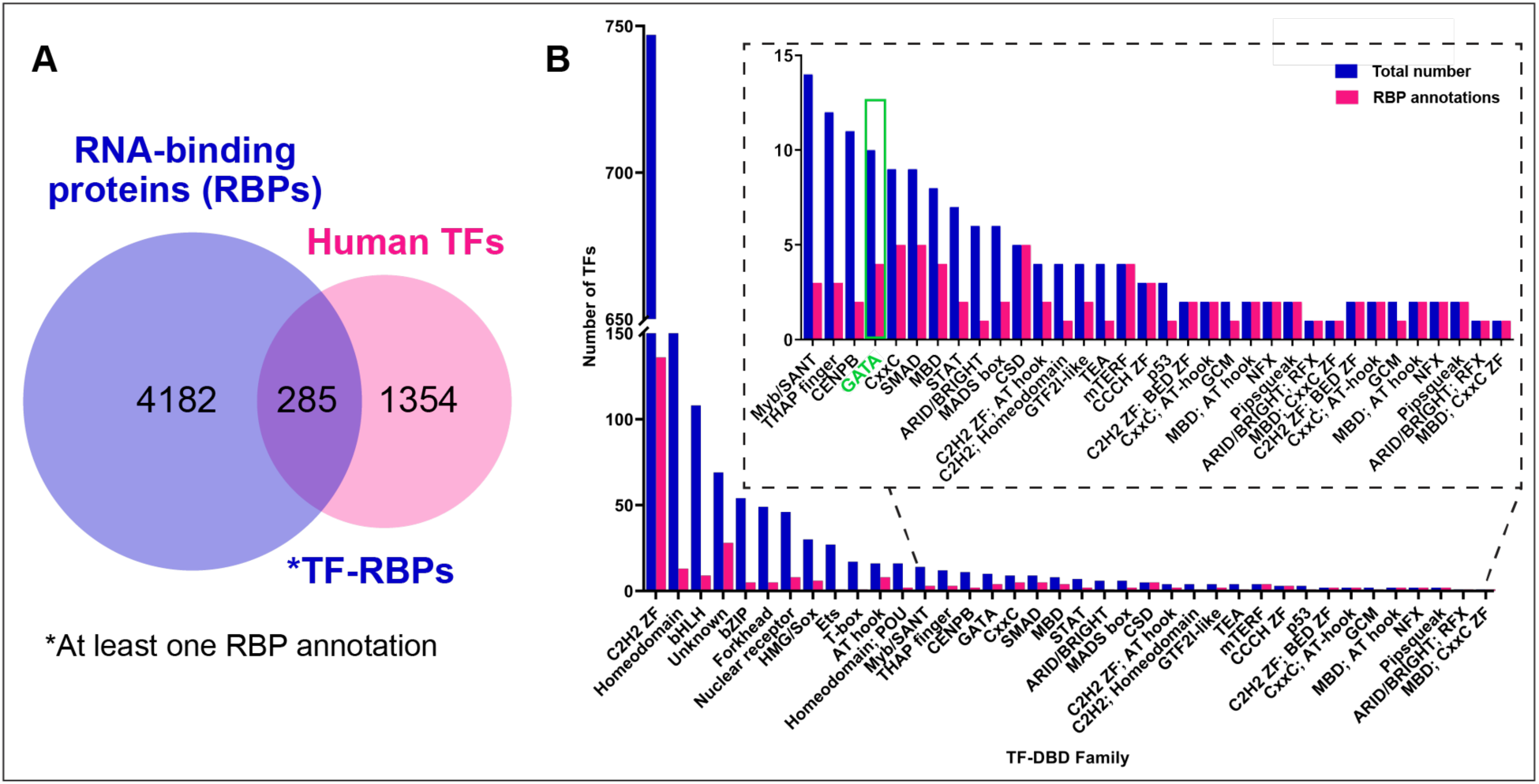
Many human TFs from various DBD families can associate with RNA. (A) Venn diagram showing an overlap for cataloged human TFs with at least one RNA-binding protein (RBP) annotation. (B) Human TFs are classified according to their DBD family with blue bars showing the total number of TFs in the respective DBD family, and magenta bars showing the number of TFs from that family with at least one RBP annotation. The inset zooms in the region of the bar chart with DBD families comprising less than 25 members.

With this dataset in hand, we next aimed to identify a model TF beyond the nuclear hormone receptor and HMG/Sox families to study competitive DNA/RNA binding. Competitive DNA/RNA binding suggests a novel mechanism for gene regulation, wherein a shared binding surface for DNA and RNA within a TF would allow the RNA to act as an immediate control switch of the TF’s ability to bind its DNA, thereby regulating transcription. To reduce the occurrence of simultaneous DNA/RNA binding TFs, we limited our search to proteins containing fewer than three DNA-binding motifs, such as zinc-fingers known to independently bind RNA,^2,32^ and surveyed literature and ENCORE datasets on selected targets. The compelling RBP candidates that emerged were members of the GATA family, which have been suggested to associate with RNA in cells.^2,3,34^ We chose to investigate the RNA-binding activity of GATA1 since it has been previously shown to crosslink with RNA in cells,^2,3^ and is a critical hemopoietic TF whose function in development and disease is a central focus of modern hematology research.^35^

### Defining DNA- and RNA-binding capacity of GATA1

To test whether GATA1 binds RNA directly, we targeted regions of the protein implicated in binding for *in vitro* testing. Examination of previously published TF-RNA-crosslinking mass spectrometry data revealed that the majority of crosslinked peptides map to the ZFs of GATA1,^3^ suggesting a potential RNA-binding surface within the DBD (Figure 2A). Prior studies by us and others reveal a trend that the RNA-binding region of TFs includes the positively charged, low-complexity regions adjacent to the DBD.^3,5,8^ Thus, we hypothesized that the RNA-binding surface of GATA1 likewise extends past the C-terminal end of the DBD to this region annotated as an ARM-like domain (Figure 2A).^3^

**Figure 2.**
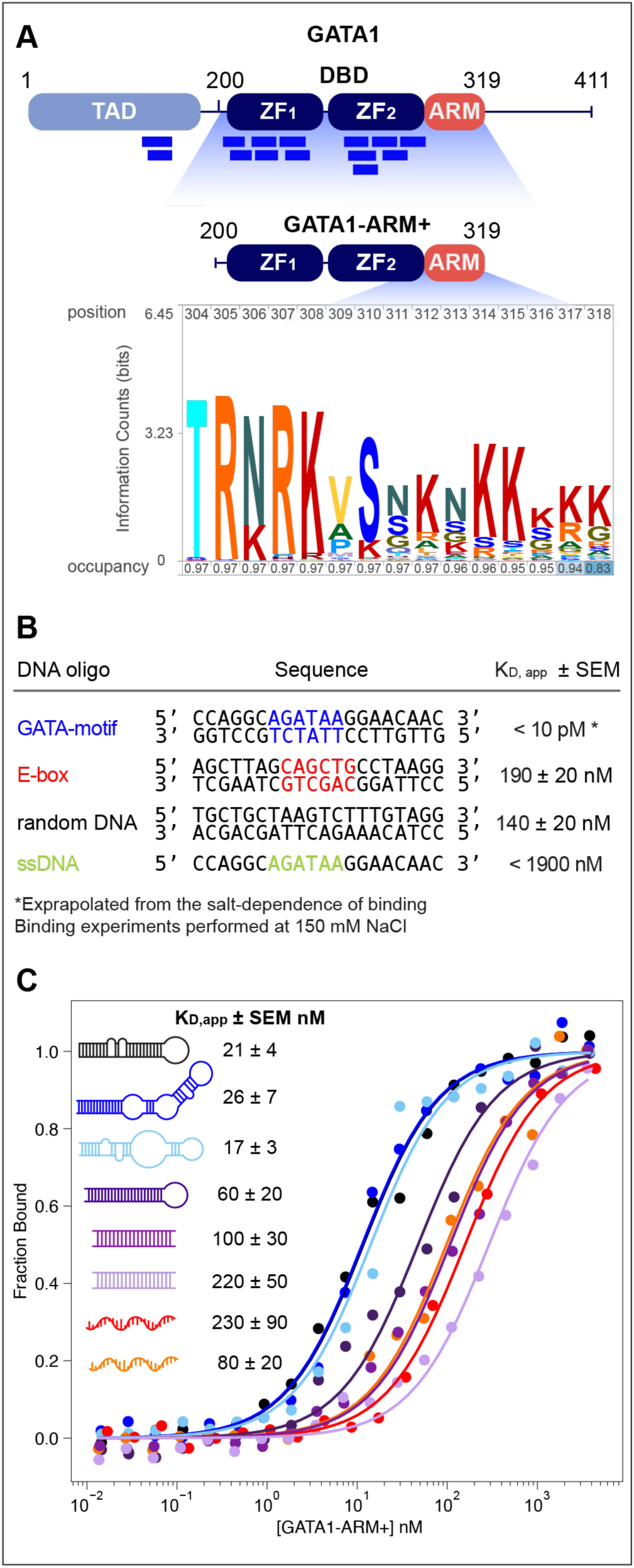
GATA1 can bind various RNA *in vitro*. (A) Domain map of full-length human GATA1, including transactivation domain (TAD), zinc fingers (ZFs) in the DBD, and the ARM-like domain (ARM). Peptides UV-crosslinked with RNA shown as blue bars were identified in the previous systematic screen of RNA-binding TFs.^3^ The recombinant construct used in this study is shown below with the peptide logo showing the conservation of the basic residues generated with Skylign.^39^ (B) and (C) binding affinities K_D, app_ and representative RNA binding isotherms (N ≥ 3, standard error of the mean reported).

To test this we recombinantly expressed and purified a construct of GATA1 containing both ZFs and ARM (GATA1-ARM+, Figure 2A) and measured its binding affinity to DNA and RNA using a fluorescence anisotropy (FA) binding assay.^36^ This construct bound GATA-motif dsDNA extremely tightly at the physiological salt concentration. Its apparent dissociation constant (K_D,app_) was estimated to a value of 10 pM based on the extrapolation from the salt-dependence of binding (Figure 2A). We note that this unusually tight binding may be attributed to our use of a full DNA-binding construct since an earlier report of sub-nanomolar binding with a dissociation constant of 0.78 nM was based on a chicken GATA1 construct containing only ZF_2_.^37^ Our construct displayed weaker binding to non-specific DNA motifs (140-190 nM) and negligible binding to ssDNA (Figure 2B), confirming high-affinity and specificity recognition of GATA-motif DNA.^37,38^

We next investigated the RNA binding and specificity of GATA1-ARM+. Since many TFs that bind RNA exhibit a preference for hairpin structures (Table S2), we probed binding to a variety of structured RNAs, including hairpin loops, hairpins with bulges, ds- and ssRNA (Figure S1). GATA1-ARM+ bound all tested RNAs with affinities ranging from 17 to 230 nM, notably tighter than non-specific DNA binding. The range of binding affinities aligned well with those observed in other TFs (Table S2). GATA1-ARM+ bound RNA hairpins and hairpins with bulges 10-fold tighter than dsRNA containing sequences that mimic DNA-binding sites such as GAUA-motif (purple) or pseudo palindromic UAUCAGAUA-motif (lavender, Figure 2C). These trends highlight a selective preference for structural elements rather than sequence-specific recognition, a common feature of many TF-RBPs.^5–8^

Unlike RNA-hairpin binding TFs such as GR, ERα, and Sox2 (Table S2),^5–8^ GATA1-ARM+ only modestly discriminates secondary structural features. The K_D,app_ for the hairpin with a terminal loop (dark purple) of 60 ± 20 nM was comparable to those for perfect duplexes (∼100-200 nM, Figure 2C). A slightly stronger binding for RNA with internal loops and unpaired bases may be explained by the reduced stiffness of the RNA duplex and, thus, more exposed surfaces for interactions with the protein. Surprisingly, GATA1-ARM+ could bind short ssRNA with GAUA-motif (230 ± 90 nM) and true ssRNA (80 ± 20 nM, Figure 2C), contrasting RNA binding specificities of GR, ERα, and Sox2.^5–8^ Taken together, these data suggest that GATA1-ARM+ is a *bona fide* RNA-binding TF with greater target promiscuity for RNA.

### GATA1 binds DNA and RNA competitively

To determine if DNA and RNA compete for binding GATA1-ARM+, we performed an equilibrium competition experiment by titrating unlabeled competitor into fluorescently labeled nucleic acid/GATA1-ARM+ complex. For this experiment, we used high-affinity nucleic acid ligands – GATA-motif dsDNA and Gas5 RNA. As expected, unlabeled GATA-motif dsDNA was able to outcompete labeled GATA-motif dsDNA from the bound complex (Figure 3A). Unlabeled GATA-motif dsDNA also competed Gas5 RNA off GATA1-ARM+, demonstrating that GATA1-ARM+ uses overlapping surfaces for RNA and DNA such that both nucleic acids cannot occupy the protein simultaneously (Figure 3B). The competitive DNA/RNA binding by GATA1 runs contrary to the proposed model for TF-DNA-RNA ternary complex.^3^ Moreover, given the high sequence and structure conservation of their DBD and ARM-like domain, this feature is likely conserved within other members of GATA family TFs (Figure 4A).

**Figure 3.**
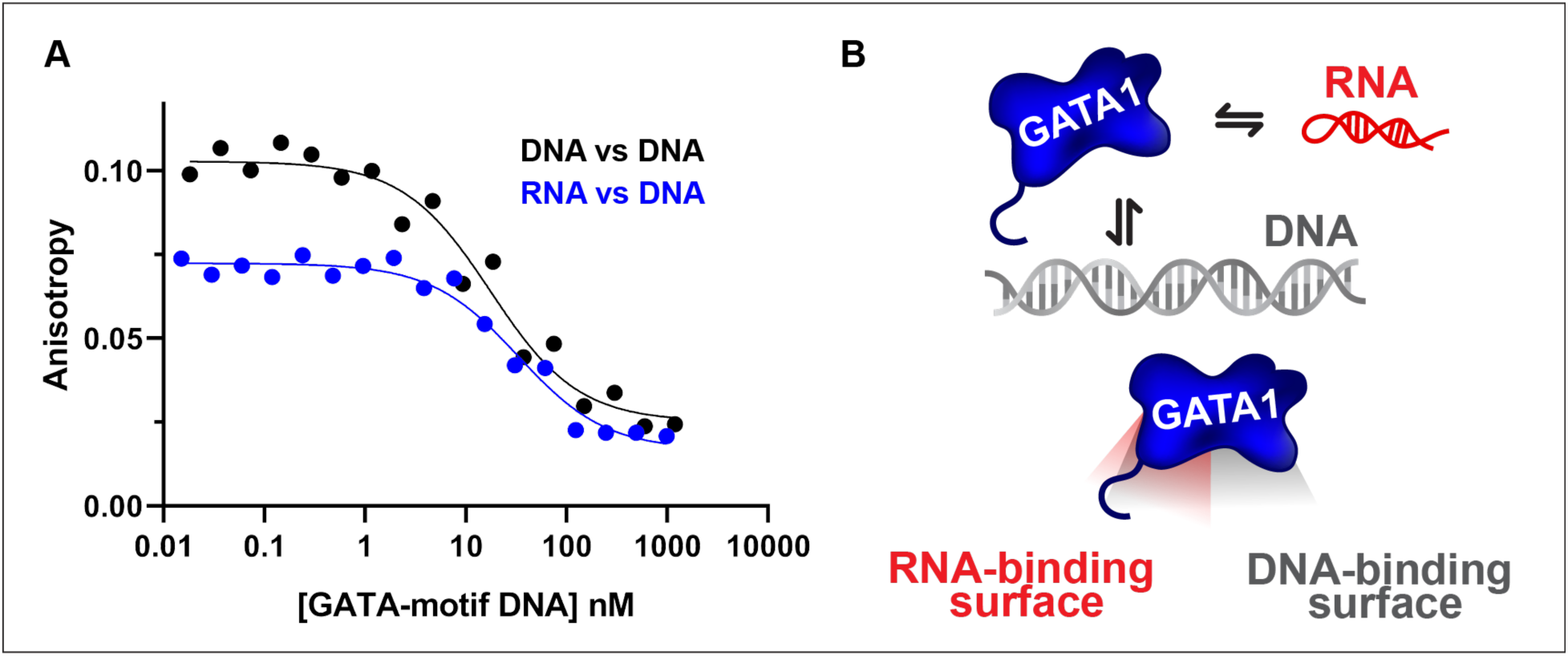
DNA and RNA binding to GATA1-ARM+ is competitive. (A) Equilibrium competition measuring the anisotropy of the labeled GATA-motif dsDNA or Gas5 RNA as a function of unlabeled GATA-motif dsDNA (N ≥ 3, standard error of the mean reported). (B) Proposed molecular model for competitive DNA and RNA binding by GATA1.

**Figure 4.**
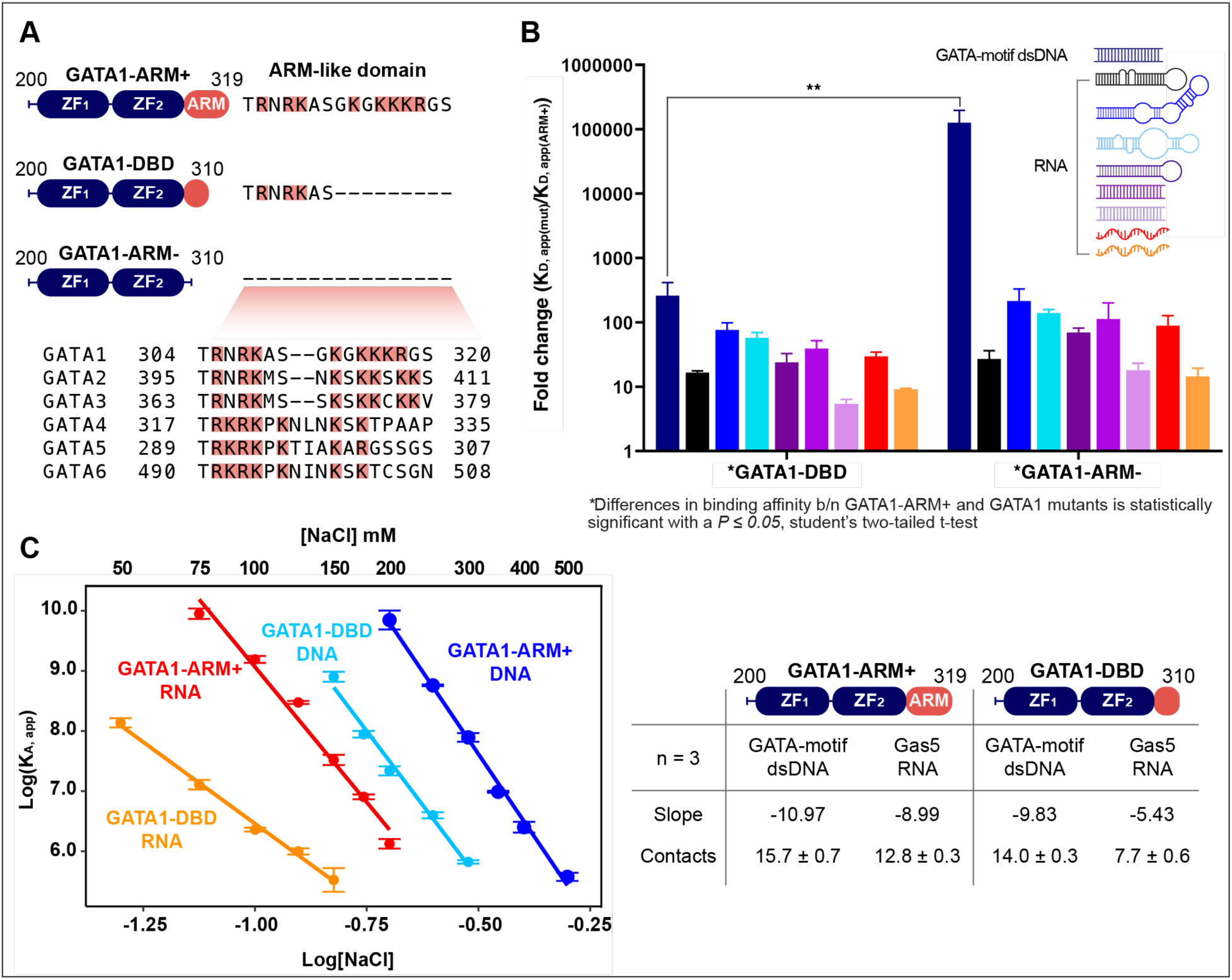
The role of the ARM-like domain in nucleic acid binding. (A) Domain map and basic patch residues of GATA1 truncation mutants; sequence alignment of the basic patch of human GATA family TFs is shown below. (B) Fold change of truncation mutants over the GATA1-ARM+ (K_D, app(mut)_/K_D, app(ARM+)_) for binding DNA (GATA-motif dsDNA) and RNA (N ≥ 3, standard error of the mean reported). (C) Double log-plot of GATA1 construct binding versus NaCl concentration for GATA-motif dsDNA and Gas5 RNA. Data were fit to the linear salt-dependence regression model.^42^ The slope and calculated values of electrostatic contacts at 150 mM NaCl are shown below.

### Redefining the role of ARM-like domain in DNA and RNA binding

The ARM-like domain, previously implicated as an independent RBD in many TFs,^3^ is well conserved across all vertebrate GATA paralogs (Figure 4A).^40^ Given GATA1’s competitive DNA/RNA binding activity, we examined how this ARM-like domain contributes to nucleic acid binding. We generated a GATA1-ARM-mutant, which lacks this domain (Figure 4A), and measured its binding affinity for GATA-motif dsDNA and RNA. Deletion of this entire region had a drastic impact on DNA binding, as the fold change of K_D, app(mut)_ over K_D, app(ARM+)_ was about 87,000 (Figure 4B). This large contribution of the ARM-like domain to DNA binding could be understood by examining the crystal structure of the DNA-bound GATA1 construct containing the N-terminal half of the ARM (PDB 3vd6).^41^ In the structure, the N-terminal portion of the ARM-like domain is an integral part of the DBD that wraps around the DNA helix in the minor groove and makes base-specific contacts via R305 and R307 (Figure S2). In distinct contrast to DNA binding, the deletion of the basic patch had a more modest impact on RNA binding (20 to 180-fold change of K_D, app(mut)_ over K_D, app(ARM+)_ our RNA ligands, Figure 4B), indicating that GATA1-RNA interactions relies on the ZFs in the DBD. These observations are consistent with competitive binding behavior, involving a shared surface for DNA and RNA rather than clearly separate domains.

We noticed the presence of two basic clusters in the ARM-like domain of GATA1 (Figure 4B), the N-terminal half belonging to the DBD and the C-terminal-half with five basic residues, which was not resolved the crystal structure (Figure S2), raising the question of whether the C-terminal half of the ARM could be preferentially involved in RNA binding over DNA binding. To test this, we generated another mutant GATA1-DBD, lacking the C-terminal half of the ARM-like domain (Figure 4A), and measured its DNA and RNA affinities. Unexpectedly, GATA1-DBD had significantly reduced DNA binding compared to the GATA1-ARM+ (155-fold change of K_D, app(mut)_ over K_D, app(ext)_, Figure 4B), emphasizing the importance of the full ARM-like domain for tight DNA binding. For RNA, the fold changes of GATA1-DBD relative to GATA1-ARM+ were comparable to those of the full ARM truncation (Figure 4B). The differences in RNA binding between GATA1-DBD and GATA-ARM-were not statistically significant and had similar trends across all tested RNA, indicating that the C-terminal part of the ARM did not considerably augment RNA binding and specificity. It appears that the entire ARM-like domain is more important for robust DNA binding by GATA1, while moderately contributing to RNA binding. Our biochemical data demonstrates that the RBD in GATA1 has been misannotated, which may also apply to other TF-RBPs.^3^

The mutagenesis data suggest that the RNA-binding surface on GATA1 includes the ZF region. To test whether electrostatic contacts relevant to RNA binding spread throughout GATA1-ARM+ and overlap with DNA-binding regions, we performed a salt-dependence assay by measuring K_D,app_ of GATA1 constructs for GATA-motif dsDNA and Gas5 RNA at varying NaCl concentrations. DNA binding by GATA1-ARM+ involved 16 electrostatic contacts, 14 of which belonged to GATA1-DBD (Figure 4C), consistent with the crystal structure.^41^ The remaining 2 contacts were mapped to the C-terminal half of the ARM. Unlike DNA binding, RNA-relevant electrostatic contacts were distributed across the whole GATA1-ARM+, with 8 contacts assigned to GATA1-DBD. These 8 RNA-relevant electrostatic contacts reinforce the involvement of DNA-binding surface in GATA1-RNA interactions and could potentially overlap with 14 DNA-relevant electrostatic contacts mapped to this region (Figure 4B). The remaining 5 electrostatic contacts for RNA binding were assigned to the C-terminal half of the ARM, corresponding to five K/R residues found in this region (Figure 4A). This exceeds the DNA-relevant contacts in the same region, indicating a stronger electrostatic contribution of the C-terminal half of the ARM to RNA binding. These interactions likely enable GATA1-ARM+ to bind diverse structured RNAs with broader specificity (Figure 2C) and explain reduced RNA affinity in GATA1-DBD (Figure 4B). The data also suggest that a simple model where the ARM-like domain is responsible for RNA binding is incorrect.^3^

## Discussion

We have identified key molecular features of GATA1-RNA interactions that point to a distinct mechanism of action. Firstly, GATA1 is a promiscuous RNA binder, whose interactome spans RNA hairpins, ds- and ssRNA fragments, far richer than that of other TF-RBPs,^3,5–8^. Secondly, GATA1 binds DNA and RNA in a mutually exclusive fashion through an extensive overlapping surface that spans the DBD and the ARM-like domain. Thirdly, the ARM-like domain contributes to DNA and RNA binding via distinct modes, which confer high-affinity DNA binding and offer more plasticity for RNA recognition driven by electrostatic interactions. We show that the presence of the basic clusters near the DNA-binding motifs does not confidently separate the DBD from a putative RBD.

Biochemical characteristics of competitive DNA/RNA binding by GATA1 may provide mechanistic insights into an RNA-centric regulation of GATA1 activity during hematopoiesis. Competitive RNA binding is expected to downregulate GATA1’s occupancy on chromatin loci that produce RNA, thus reducing GATA1 transcriptional activity.^10^ Considering the essential role of GATA1 in erythroid and megakaryocytic differentiation,^35^ GATA1-RNA interactions could modulate GATA1 occupancy on lineage-sensitive loci and direct transcriptional programs toward a specific lineage during hematopoiesis. It is also possible that RNA binding could affect chromatin looping by directly competing with tethering DNA contacts since GATA1 has been shown to establish long-distance promoter-enhancer interactions via LDB1 and cohesion complex during erythropoiesis.^43^ Functional *in vivo* studies are needed to establish the role of GATA1-RNA interactions during hematopoiesis.

Our study emphasizes the diversity of binding mechanisms of non-canonical RBPs as well as the importance of a robust biochemical approach for dissecting these interactions. The expanding set of competitive DNA/RNA-binding TFs highlights the unappreciated multifunctionality of DBDs in nucleic acid recognition, challenging the prevailing idea of functionally separate binding domains. While the presence of motifs resembling RBDs such as ARM-like domain could indicate RNA-binding activity,^3^ these motifs may work in concert with DBDs to offer dynamic interplay between DNA and RNA binding, as in the case of GATA1. Molecular signatures of RNA binding by TFs may match the diversity of DBDs implicated in RNA interactions (Figure 1B). Therefore, delineating DBDs and non-canonical RBDs may become particularly challenging when screening newly discovered DNA/RNA binding proteins *en masse* using predictive models without rigorous biochemical characterization.

## Supporting information

supplement_gata

## ASSOCIATED CONTENT

### Supporting Information

Materials and methods, supplementary tables, and figures (PDF)

Table S1. RNA-interacting transcription factors (TableS1_TFRBP.xlsx)

GATA1 – UniProtKB P15976

## AUTHOR INFORMATION

### Author Contributions

D.AU., R.T.B., and D.S.W. designed the research. D.A.U. conducted experimental research and wrote the paper with feedback from all authors, R.T.B. and D.S.W. supervised the project.

## ACKNOWLEDGMENT

We thank Dr. Nickolaus Lammer for helping with bioinformatic search of RNA-binding TFs and the Shared Instruments Pool facility (RRID: SCR_018986, NIH Grant R24OD033699-01, NIH Shared Instrumentation Grant S10OD034218-0) in the Department of Biochemistry, University of Colorado Boulder for the use of shared instruments.

## ABBREVIATIONS

TF: transcription factor;
RBP: RNA-binding protein;
DBD: DNA-binding domain;
RBD: RNA-binding domain;
GR: glucocorticoid receptor;
ERα: estrogen receptor alpha,
FA: fluorescence anisotropy.

