## supplement_gata for "A distinct mechanism of RNA recognition by the transcription factor GATA1": 02_supplementary_bioarxiv.pdf

### Materials and Methods

#### Analysis of RNA-binding transcription factors

A list of previously documented RNA-interacting proteins was obtained R-Deep website that curates RNA-binding and RNA-dependent proteins from twenty independent studies.[1–20] This dataset was combined with a curated catalog of known and likely human transcription factors (TFs)[21] by finding matched proteins in both datasets using their gene identifiers (HGNC ID and gene names) in RStudio. The matched hits were identified as putative RNA-interacting TFs if they had at least one RNA-binding protein annotation. These RNA-interacting TFs were sorted according to their respective DNA-binding domain (DBD) family.[21] The full list of these RNA-interacting TFs along with the respective DBD family and total number of RBP annotations can be found in TableS1_TFRBP.xlsx.

#### Cloning, Expression, and Purification of recombinant GATA1 constructs

Human GATA1 gene was obtained from Addgene in a lentiviral vector (Plasmid #61062). GATA1-ARM+ region (residues 200-319) was cloned into a pET28b expression vector containing N-terminal 10xHis-SUMO tag via Gibson assembly. GATA1-DBD and GATA1-ARM- constructs were generated by site-directed mutagenesis using mutagenic primers. All Gibson assembled plasmids were verified by Nanopore sequencing (Quintara Biosciences). Protein sequences of GATA1 constructs used in this study are listed in Table S2.

Recombinant proteins were expressed in *Escherichia coli* BL21(DE3) pLysS expression cells. A starter culture was made by growing a single transformed colony in 40 mL LB media with 50 µg/mL kanamycin and 34 µg/mL chloramphenicol overnight at 37°C with continuous shaking. Expression cultures were prepared by inoculating 10 mL of the overnight starter in 1L of 2x YT media containing 50 µg/mL kanamycin and 34 µg/mL chloramphenicol. The cultures were grown at 37°C to $OD_{600}$ of 0.8-1.0 followed by a 30 min incubation on ice. Protein expression was induced with 0.5 mM isopropyl-β-d-thiogalactopyranoside (IPTG) and 50 µM $ZnCl_2$ at 37°C with continuous shaking for 3 hrs. Cells were harvested by centrifugation at 5,000 RCF at 4°C for 10 min (Avanti JXN-26 floor centrifuge). The pellets were stored at -20°C.

All protein constructs were subjected to the same 3-column purification protocol. Cell pellet (3 L expression) was resuspended in 80 mL ice-cold lysis buffer (20 mM Tris pH 7.5, 0.5 M NaCl, 10 mM imidazole pH 7.5, 10% glycerol) with two cOmplete EDTA-free EASYpack protease inhibitor tablet (Roche) tablets using 100-mL tissue Dounce homogenizer. Cells were lysed via sonication on ice with a QSonica Q500 Sonicator for 7 min in pulses of 15 s followed by 45 s of rest. The lysate was cleared by centrifugation at 27,000 RCF at 4°C for 30 min (Avanti JXN-26 floor centrifuge) and incubated with 16 mL precleared Ni/NTA beads at 4°C for 1 hr, after which the unbound protein was released by gravity flow through the column. The beads were washed three times – 1) lysis buffer; 2) lysis buffer containing 30 mM imidazole; 3) lysis buffer containing 50 mM imidazole. The protein was eluted with 100 mL lysis buffer containing 350 mM imidazole, and the eluate was concentrated down to 50 mL in 10 kDa MWCO Amicon® concentrator (Millipore Sigma). Every 10 min throughout the concentration process, the sample was resuspended by gentle pipetting to prevent aggregation on the regenerated cellulose membrane. The final protein concentrate was filtered through a cellulose acetate membrane filter with 0.2 micron pore size (Advantec 25CS020AS), transferred into 6–8K MWCO dialysis tubing (Spectra/Por – Spectrum Labs), and dialyzed at 4°C in the buffer containing 20 mM Tris pH 7.5, 300 mM NaCl, 5% glycerol, and 1 mM DTT. After 2.5-3 h of dialysis, one mg of in-house His-tagged ULP1 SUMO-protease was added to the partially dialyzed protein sample to cleave the 10xHis-SUMO tag, and the dialysis proceeded overnight at 4°C. Dialyzed protein sample was filtered through a cellulose acetate membrane filter with 0.2 micron pore size (Advantec 25CS020AS) before loading onto 50 mL superloop for purification through 5 mL HisTrap HP column (Cytiva) pre-equilibrated with buffer A (20 mM Tris pH 7.5, 300 mM NaCl, 5% glycerol, and 1 mM DTT) and 2% buffer B (500 mM imidazole pH 7.5, 20 mM Tris pH 7.5, 300 mM NaCl, 5% glycerol, and 1 mM DTT). Following the sample application, the column was washed twice with 10 column volumes – once with 10% buffer B, and once with 20% buffer B. Untagged protein was collected in the second wash with 20% buffer B, and the tag was eluted with 30-100% buffer B linear gradient. Eluted untagged protein was concentrated to 2 mL in 3 kDa MWCO Amicon® concentrator (Millipore Sigma) with continuous mixing every 10 min to prevent aggregation on the membrane. The concentrated sample was filtered in Spin-X® cellulose acetate centrifuge tubes (Costar) prior to injection onto Superdex™ 75 Increase 5/150 GL column (Cytiva) pre-equilibrated in 20 mM Tris pH 7.5, 300 mM NaCl, 5% glycerol, 1 mM DTT buffer. Fractions containing the purified protein were pooled, concentrated to 100-120 µM in a 3 kDa MWCO Amicon® concentrator (Millipore Sigma) with mixing every 3 min, filtered in Spin-X® cellulose acetate centrifuge tubes, aliquoted, flash frozen, and stored at −70°C. The protein concentration was determined via UV-Vis spectroscopy by measuring the absorbance at 280 nm and using the extinction coefficient of 16,960 $M^{-1}$ $cm^{-1}$ for all GATA1 contracts used in this study. The typical yield ranged between 0.8 to 1.5 mg/L of growth.

**Oligonucleotide preparation**
Oligonucleotide sequences used in this study are listed in Table S2. All double-stranded (ds) DNA oligos were generated by annealing single-stranded (ss) DNA oligos. Briefly, ssDNA oligos were obtained from Integrated DNA Technologies (IDT) with standard desalting purification, the sense strands were conjugated with 6-FAM at the 5'-end. ssDNA oligos were annealed in 20 mM Tris pH 7.5, 50 mM NaCl buffer at 1 or 2 µM by boiling at 95°C for 2 min and slow cooling to room temperature for 3 h. Annealed dsDNA oligos were stored at 4°C.

dsRNA oligos were generated by annealing ssRNA oligos obtained from IDT. The forward strands were conjugated with 6-FAM at the 5'-end. Annealed reactions were made in 20 mM Tris pH 7.5, and 10 mM NaCl buffer at 1 or 2 µM by boiling at 95°C for 2 min and slow cooling to room temperature for 3 h. Annealed dsRNA oligos were stored at 4°C until use.

Structured RNA oligos were prepared by *in vitro* transcription using T7 RNA polymerase and dsDNA templates. DNA templates with T7 promoter sequence at the 5'-end were obtained from IDT and PCR amplified using a forward primer (T7 promoter sequence) and reverse primer unique to the 3'-end of each template. *In vitro* transcription reactions were performed as described in previously published protocols,[22] and RNA oligos were purified by denaturing gel electrophoresis (1x TBE, 8 M urea). Purified RNAs were labeled via 3'-end ligation of pCp-ATTO-488 (Jena Bioscience) by T4 ssRNA Ligase (NEB) as described previously.[23,24] RNA concentration was determined via UV-Vis spectroscopy by measuring the absorbance at 260 nm and using an extinction coefficient of each RNA (inferred from sequence). The typical RNA yield from the labeling reaction ranged between 40-60% with 70-80% labeling efficiency. RNA purity and labeling were determined by running a denaturing PAGE (15% polyacrylamide, 1x TBE, 8 M urea) and imaging for the label using Typhoon FLA 9500. Fluorescently labeled RNAs were stored at -20°C until use. Before setting up fluorescence anisotropy (FA) binding reactions, fluorescently labeled RNA ligands were diluted to 1 µM in refolding buffer (20 mM Tris pH 7.5, 10 mM NaCl) and refolded by boiling at 95°C for 1 min and cooling on ice for 5 min.

**Fluorescent anisotropy binding assay**
The assay was set up based on previously published protocols.[23,25,26] RNA binding reactions were prepared in the binding buffer containing 20 mM Tris pH 7.5, 150 mM NaCl, 1 mM DTT, 0.01% IGEPAL). Typical DNA concentrations for strong binding (e.g., GATA1-DBD construct and GATA-motif dsDNA) were held constant at 0.35 nM in the final binding reaction to maintain good signal-to-noise. Typical RNA ligand concentration was held at 1 to 2 nM in the final binding reaction. To perform equilibrium binding, purified protein was serially diluted to 2x final binding concentration in the binding buffer. Protein titrations typically ranged from 10-5 µM to 0.015 nM.

DNA or RNA ligands were separately diluted at 2x final binding concentration. Protein and ligand were mixed in a 1:1 volume ratio in a 20 μL binding reaction in a flat-bottom low-flange 384-well black NBS polystyrene plate (Corning) and were allowed to reach equilibrium at room temperature in the dark for 1 hr. Parallel ($I_{\parallel}$) and perpendicular ($I_{\perp}$) fluorescence intensities were measured with ClarioStar Plus FP plate reader (BMG Labtech), and anisotropy for each protein dilution point was calculated using $Anisotropy = (I_{\parallel} + I_{\perp})/(I_{\parallel} + 2 \times I_{\perp})$ . Anisotropy values were plotted as a function of the log of protein concentration and corresponded to 1:1 binding stoichiometry for both DNA and RNA binding. The apparent dissociation constant, $K_{D,app}$, was determined by fitting the data to the quadratic binding isotherm

$$Anisotropy = S\left(\frac{[L]_T+[P]_T+K_{D,app}-\sqrt{([L]_T+[P]_T+K_{D,app})^2-4([L]_T[P]_T)}}{2[L]_T}\right)+O,$$

where S is the difference between the max and min anisotropy values, O is an offset factor equivalent to the min anisotropy value, and $[P]_T$ and $[L]_T$ are known concentrations of GATA1 and DNA/RNA, respectively. The data was fit using in-house generated script (https://github.com/nicklammer/AnisotropyBindingFit). Each binding reaction was performed at least in triplicate using different protein and ligand dilutions on separate days and remained consistent across independent protein and RNA preparations. The statistical significance was determined for differences between average values of $K_{D,app}$ using student's two-tailed t-test.

**Critical considerations for FA-based assays**

An important consideration in fluorescence-based binding experiments is monitoring the change in total fluorescence intensity of the probe upon protein binding. The total fluorescence intensity is calculated from parallel and perpendicular intensities

$$I_{tot} = I_{\parallel} + 2I_{\perp},$$

where $I_{tot}$ is the total fluorescence intensity, $I_{\parallel}$ is the parallel intensity, and $I_{\perp}$ is the perpendicular intensity.

In our DNA binding experiments, the binding was accompanied by no significant changes in total fluorescence intensity upon protein binding (Figure S4). For RNA binding tested with in-house 3'- ATTO-488 labeled RNA, the total fluorescence intensity was quenched with increasing protein concentrations. We note that the apparent dissociation constants calculated using the quenching fit were not significantly different from the anisotropy binding fit calculations (Figure S5).

$$Quenching = \frac{I_0-I_{obs}}{I_0} = Q_{max}\left(\frac{K_A[P]}{1+K_A[P]}\right),$$ [27]

Where $I_0$ is the total fluorescence intensity of the free probe, $I_{obs}$ is the total fluorescence intensity observed at a given protein concentration, $Q_{max}$ is the maximum observed quenching, $K_A$ is the association constant, [P] is the protein concentration.

To ensure that the label and its position do not affect the measured binding affinity of the protein for RNA, we performed FA binding assay with GATA1-ARM+ construct and Gas5 RNA with 6-FAM label at the 5'-end (RNA obtained from Horizon Discovery, Dharmacon). The binding was performed in the same conditions as described in the previous section. The apparent dissociation constants for 5'-6FAM and in-house 3'- ATTO-488 labeled Gas5 RNA were very similar (Figure S6). Thus, we proceeded with measuring the RNA binding by GATA1 using in-house 3'-ATTO-488 labeled RNA.

**Competition and salt-dependence experiments**

Equilibrium competition experiments were performed in the same binding buffer (20 mM Tris pH 7.5, 150 mM NaCl, 1 mM DTT, 0.01% IGEPAL)) by measuring fluorescence anisotropy. Fluorescently labeled ligand and purified GATA1 were mixed such that the ligand was bound by the protein at 70% followed by titration of an unlabeled competitor. Typical competitor concentrations varied between 100-fold below and 100-fold $K_{D,app}$. The competition reactions were equilibrated in the dark for 1 hr, and intensities were measured with ClarioStar Plus FP plate reader (BMG Labtech). Anisotropy values were calculated as described above and plotted as a function of competitor concentration. The data were fit in Prism (GraphPad) using a dose-response-inhibition model for IC50 determination and one site competitive binding model for $K_I$ determination. Each competition experiment was performed at least in triplicate with independent protein, ligand, and competitor dilutions. The competition experiments remained consistent across independent protein and RNA preparations.

Salt-dependence of binding for GATA-motif dsDNA and Gas5 RNA was performed according to the previously published studies.[25,28] The apparent dissociation constants were determined by measuring anisotropy in the binding buffer with varying NaCl concentrations (20 mM Tris pH 7.5, 1mM DTT, 0.01% IGEPAL, and NaCl ranging from 50 to 500 mM). The double logarithmic plots were fit to the linear regression model

$$Log\left(K_{A,app}\right) = Log\left(K_{A,nel}\right) - N \times Log[NaCl],$$

where the slope of the line $N = 0.7 \times Z$, and $Z$ is the number of electrostatic contacts, $Log\left(K_{A,nel}\right)$ is the non-electrostatic component. The underlying assumptions and limitations of the model are described in previous reports.[29,30]

**Table S1. RNA-interacting transcription factors**
TableS1_TFRBP.xlsx

**Table S2. Binding affinities of validated RNA-binding transcription factors**

| TF | TF-DBD Family | RNA ligands | $K_D$ range | References |
|---|---|---|---|---|
| GATA1 | GATA | Short hairpins, double- and single-stranded RNA | 20-230 nM | This study |
| GR | Nuclear receptor | Short hairpins | 30-70 nM | Parsonnet et al.[25] |
| ERα | Nuclear receptor | Short hairpins, hairpins with internal loops | 40-180 nM | Steiner et al.[23] |
| Sox TFs | HMGB-Sox | ES2 lncRNA, stem-loops with intranal loops, RNA 4-way junction, TERRA rG4 | < 1 nM to 3 μM | Holmes et al., Hamilton et al.[28,31] |
| TFIIIA | $C_2H_2$ zinc finger | 5S rRNA | 1-2 nM | Clemens et al.[32] |
| CTCF | $C_2H_2$ zinc finger | Tsix, Xite, Jpx lncRNA | < 1 nM | Kung et al.[33] |
| YY1 | $C_2H_2$ zinc finger | SELEX-generated RNA | 4 μM | Wai et al.[34] |
| Smad3 | Smad | Longer hairpins with internal loops and junctions | 190-570 nM | Dickey et al.[35] |

**Table S3. GATA1 recombinant protein information (GATA1 – UniProtKB P15976)**

| Name | Residues | Sequence | Size, kDa | pI |
|---|---|---|---|---|
| 10xHis-SUMO-GATA1-ARM+ | 200-319 | MGHHHHHHHHHHSSGHIEGRHMASMSDSEVNQEAKPEVKPEVKPETHINLKVSDGSSEIFFKIKKTTPLRRLMEAFAKRQGKEMDSLRFLYDGIRIQADQTPEDLDMEDNDIIEAHREQIGGSEARECVNCGATATPLWRRDRTGHYLCNACGLYHKMNGQNRPLIRPKKRLIVSKRAGTQCTNCQTTTTTLWRRNASGDPVCNACGLYYKLHQVNRPLTMRKDGIQTRNRKASGKGKKKRGS | 27.7 | 9.63 |
| GATA1-ARM+ | 200-319 | EARECVNCGATATPLWRRDRTGHYLCNACGLYHKMNGQNRPLIRPKKRLIVSKRAGTQCTNCQTTTTTLWRRNASGDPVCNACGLYYKLHQVNRPLTMRKDGIQTRNRKAS | 13.6 | 10.55 |
| 10xHis-SUMO-GATA1-DBD | 200-310 | MGHHHHHHHHHHSSGHIEGRHMASMSDSEVNQEAKPEVKPEVKPETHINLKVSDGSSEIFFKIKKTTPLRRLMEAFAKRQGKEMDSLRFLYDGIRIQADQTPEDLDMEDNDIIEAHREQIGGSEARECVNCGATATPLWRRDRTGHYLCNACGLYHKMNGQNRPLIRPKKRLIVSKRAGTQCTNCQTTTTTLWRRNASGDPVCNACGLYYKLHQVNRPLTMRKDGIQTRNRKAS | 26.8 | 9.31 |
| GATA1-DBD | 200-310 | EARECVNCGATATPLWRRDRTGHYLCNACGLYHKMNGQNRPLIRPKKRLIVSKRAGTQCTNCQTTTTTLWRRNASGDPVCNACGLYYKLHQVNRPLTMRKDGIQTRNRKAS | 12.6 | 10.27 |
| 10xHis-SUMO-GATA1-ARM- | 200-303 | MGHHHHHHHHHHSSGHIEGRHMASMSDSEVNQEAKPEVKPEVKPETHINLKVSDGSSEIFFKIKKTTPLRRLMEAFAKRQGKEMDSLRFLYDGIRIQADQTPEDLDMEDNDIIEAHREQIGGSEARECVNCGATATPLWRRDRTGHYLCNACGLYHKMNGQNRPLIRPKKRLIVSKRAGTQCTNCQTTTTTLWRRNASGDPVCNACGLYYKLHQVNRPLTMRKDGIQ | 25.9 | 9.02 |
| GATA1-ARM- | 200-303 | EARECVNCGATATPLWRRDRTGHYLCNACGLYHKMNGQNRPLIRPKKRLIVSKRAGTQCTNCQTTTTTLWRRNASGDPVCNACGLYYKLHQVNRPLTMRKDGIQ | 11.8 | 9.95 |

**Table S4. DNA and RNA oligo information (all oligos are color-coded as they appear in the main text)**

| Features | Name | Sequence 5'-3' | Label | Size (bp/nt) |
|---|---|---|---|---|
| dsDNA | GATA-motif | CCAGGCAGATAAGGAACAAC | 5'-6FAM | 20 |
| dsDNA | E-box | AGCTTAGCAGCTGCCTAAGG | 5'-6FAM | 20 |
| dsDNA | random DNA | TGCTGCTAAGTCTTTGTAGG | 5'-6FAM | 20 |
| ssDNA | GATA-motif FWD strand | CCAGGCAGATAAGGAACAAC | 5'-6FAM | 20 |
| structured | Gas5 | GGUGCCUCCCAGUGGUCUUUGUAG ACUGCCUGAUGGAGUCACCC | 3'-AF488 | 45 |
| structured | Ccl2 | GGAGAGGCUGAGACUAACCCAGAAA CAUCCAAUUCUCAAACUGAAGCUCG CACUCUCCC | 3'-AF488 | 59 |
| structured | 7SK-GAGA | GGGAUCUGUCACCCCAUUGAUCGC CGAGAGGCUGAUCUGGCUGGCUAG GCGGGUCCCC | 3'-AF488 | 58 |
| structured | Gas5 hairpin | GGUGCCUCCAUCACCUGGUCUUUG UAGACCACCUGAUGGAGGCACCC | 3'-AF488 | 48 |
| dsRNA | rGATA | CCAGGCAGAUAAGGAACAAC | 5'-6FAM | 20 |
| dsRNA | rmPal | AGUCCAUCUGAUAAGACUUCA | 5'-6FAM | 21 |
| ssRNA | env8 short | AUACAACAUACAACAUACAA | 5'-6FAM | 20 |
| ssRNA | rGATA FWD strand | CCAGGCAGAUAAGGAACAAC | 5'-6FAM | 20 |

**Table S5. Binding affinity constants for GATA1 constructs used in this study. All binding experiments were performed in 150 mM NaCl binding buffer at least in triplicate.**

| Construct | DNA/RNA ligand | $K_{D,app}$ ± SEM | Fold Change ($K_{D,app\,(mut)}/K_{D,app(ARM+)}$) |
|---|---|---|---|
| GATA1-ARM+ | dsGATA-motif | < 10 pM* | - |
| | ssDNA | < 1900 nM | - |
| | dsE-box | 190 ± 20 nM | - |
| | random dsDNA | 140 ± 20 nM | - |
| GATA1-DBD | dsGATA-motif | 1.2 ± 0.2 nM | 150 |
| | ssDNA | < 11,000 nM | 6 |
| | dsE-box | < 2300 nM | 12 |
| | random dsDNA | < 1500 nM | 11 |
| GATA1-ARM- | dsGATA-motif | 670 ± 150 nM | 87,000 |
| GATA1-ARM+ | Gas5 | 21 ± 4 nM | - |
| | Gas5 hairpin | 60 ± 20 nM | - |
| | 7SK-GAGA | 26 ± 7 nM | - |
| | Ccl2 | 17 ± 3 nM | - |
| | rGATA dsRNA | 100 ± 30 nM | - |
| | rmPal dsRNA | 220 ± 50 nM | - |
| | env8 | 230 ± 90 nM | - |
| | rGATA ssRNA | 80 ± 20 nM | - |
| GATA1-DBD | Gas5 | 510 ± 70 nM | 24 |
| | Gas5 hairpin | < 1300 nM | 21 |
| | 7SK-GAGA | < 1400 nM | 57 |
| | Ccl2 | < 1000 nM | 64 |
| | rGATA dsRNA | <2300 nM | 23 |
| | rmPal dsRNA | <1400 nM | 6 |
| | env8 | <8600 nM | 38 |
| | rGATA ssRNA | <1800 nM | 22 |
| GATA1-ARM- | Gas5 | 780 ± 80 nM | 36 |
| | Gas5 hairpin | < 2800 nM | 46 |
| | 7SK-GAGA | < 3200 nM | 123 |
| | Ccl2 | < 2900 nM | 178 |
| | rGATA dsRNA | < 5000 nM | 51 |
| | rmPal dsRNA | < 5000 nM | 23 |
| | env8 | < 11,000 nM | 52 |
| | rGATA ssRNA | < 2100 nM | 26 |

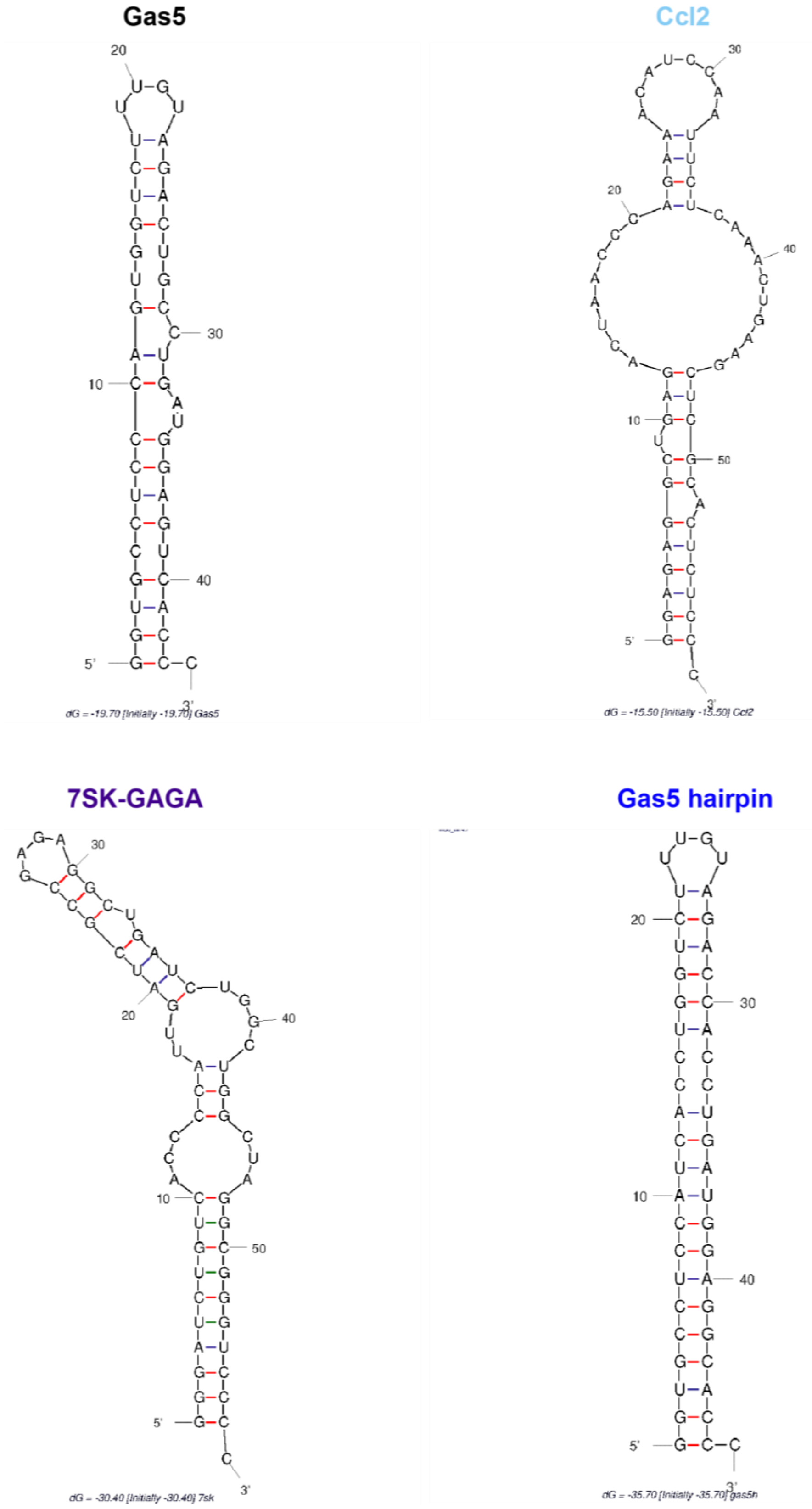


**Figure S1**. Secondary structures of RNA ligands used in this study. Structures were predicted by UNAFold.[36]

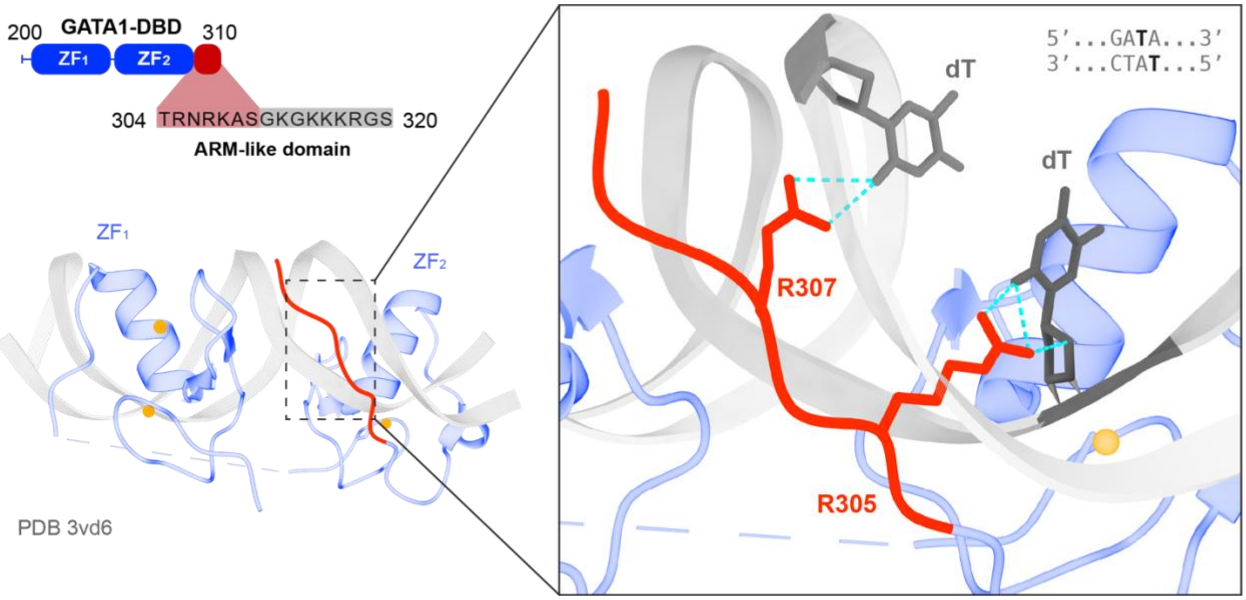


**Figure S2**. Crystal structure of GATA1-DBD bound to murine pseudo palindromic GATA-motif containing DNA. The domain map of GATA1-DBD is shown on the top, the C-terminal end of the ARM-like domain (grey) is not resolved in the crystal structure. The inset shows base-specific interactions established by R305 and R307 residues with the GATA-motif, hydrogen bonds are shown as dashed cyan lines.

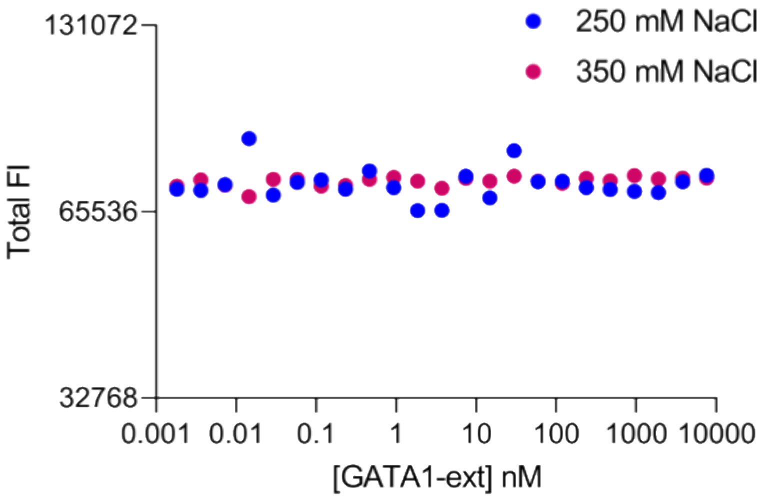


**Figure S3**. The total fluorescence intensity (FI) of 5'-6FAM labeled GATA-motif dsDNA monitored across varying GATA1-ARM+ concentrations in the course of DNA binding in 250 mM or 350 mM NaCl containing binding buffer.

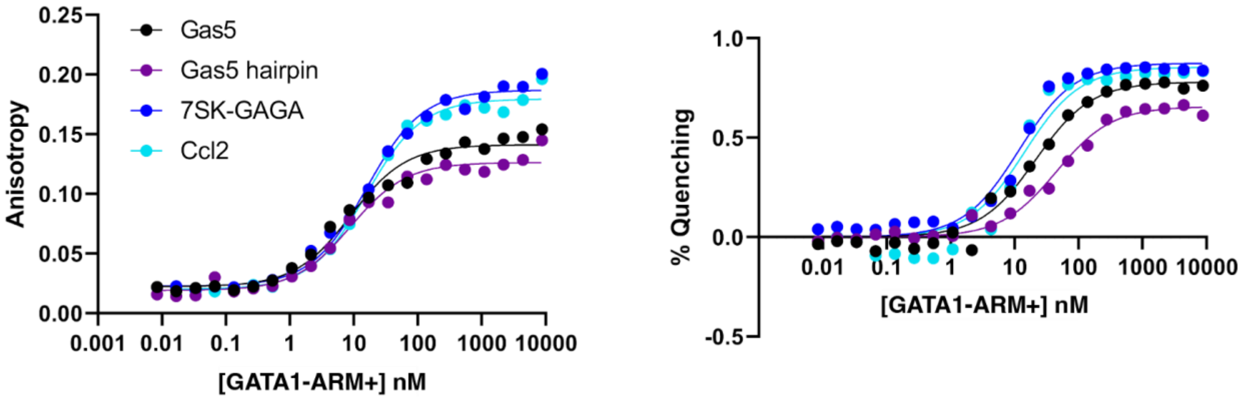


| RNA | $K_{D, app}$ nM (n = 3) | |
|---|---|---|
| | Anisotropy fit | Quenching fit |
| Gas5 | 8.2 ± 1.7 | 15.9 ± 6.6 |
| Gas5 hairpin | 7.7 ± 1.1 | 30.8 ± 13.6 |
| 7SK | 12.6 ± 3.8 | 8.6 ± 2.3 |
| Ccl2 | 13.2 ± 2.0 | 10.8 ± 2.4 |

**Figure S4**. Representative binding isotherms for GATA1-ARM+ and in-house 3'-ATTO-488 labeled RNA were analyzed using anisotropy fit (left) and quenching fit (right). The values for apparent dissociation constants $K_{D, app}$ are listed in the table below.

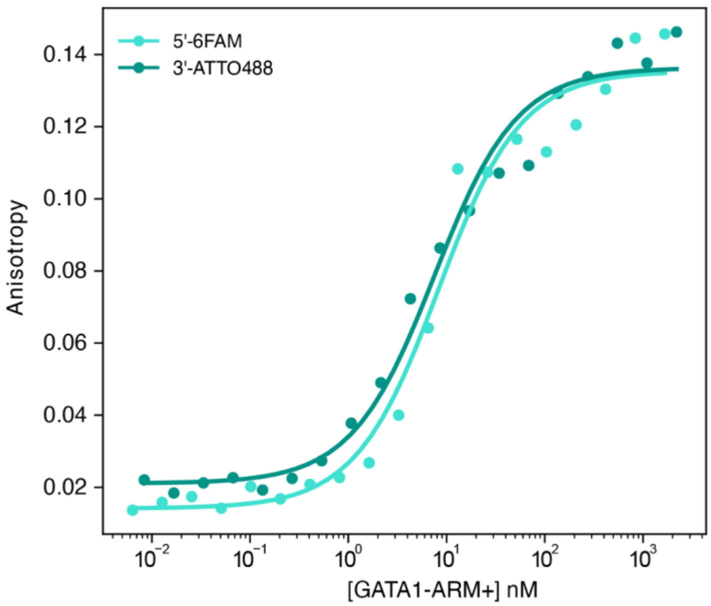


| n = 3 | 5'-6FAM | 3'-ATTO488 |
|---|---|---|
| $K_{D,app}$ nM | 9.7 ± 0.4 | 8.2 ± 1.7 |

**Figure S6**. Representative binding isotherms comparing GATA1-ARM+ binding to 5'-FAM labeled Gas5 and in-house 3'-ATTO-488 labeled Gas5. The values for apparent dissociation constants $K_{D,\ app}$ calculated using anisotropy fit are listed in the table.

(13) Ray, D.; Kazan, H.; Cook, K. B.; Weirauch, M. T.; Najafabadi, H. S.; Li, X.; Gueroussov, S.; Albu, M.; Zheng, H.; Yang, A.; Na, H.; Irimia, M.; Matzat, L. H.; Dale, R. K.; Smith, S. A.; Yarosh, C. A.; Kelly, S. M.; Nabet, B.; Mecenas, D.; Li, W.; Laishram, R. S.; Qiao, M.; Lipshitz, H. D.; Piano, F.; Corbett, A. H.; Carstens, R. P.; Frey, B. J.; Anderson, R. A.;

Lynch, K. W.; Penalva, L. O. F.; Lei, E. P.; Fraser, A. G.; Blencowe, B. J.; Morris, Q. D.; Hughes, T. R. A Compendium of RNA-Binding Motifs for Decoding Gene Regulation. *Nature* **2013**, *499* (7457), 172–177. https://doi.org/10.1038/nature12311.
(14) Gerstberger, S.; Hafner, M.; Tuschl, T. A Census of Human RNA-Binding Proteins. *Nat Rev Genet* **2014**, *15* (12), 829–845. https://doi.org/10.1038/nrg3813.
(15) He, C.; Sidoli, S.; Warneford-Thomson, R.; Tatomer, D. C.; Wilusz, J. E.; Garcia, B. A.; Bonasio, R. High-Resolution Mapping of RNA-Binding Regions in the Nuclear Proteome of Embryonic Stem Cells. *Molecular Cell* **2016**, *64* (2), 416–430. https://doi.org/10.1016/j.molcel.2016.09.034.
(16) G Hendrickson, D.; Kelley, D. R.; Tenen, D.; Bernstein, B.; Rinn, J. L. Widespread RNA Binding by Chromatin-Associated Proteins. *Genome Biology* **2016**, *17* (1), 28. https://doi.org/10.1186/s13059-016-0878-3.
(17) Sundararaman, B.; Zhan, L.; Blue, S. M.; Stanton, R.; Elkins, K.; Olson, S.; Wei, X.; Van Nostrand, E. L.; Pratt, G. A.; Huelga, S. C.; Smalec, B. M.; Wang, X.; Hong, E. L.; Davidson, J. M.; Lécuyer, E.; Graveley, B. R.; Yeo, G. W. Resources for the Comprehensive Discovery of Functional RNA Elements. *Molecular Cell* **2016**, *61* (6), 903–913. https://doi.org/10.1016/j.molcel.2016.02.012.
(18) Brannan, K. W.; Jin, W.; Huelga, S. C.; Banks, C. A. S.; Gilmore, J. M.; Florens, L.; Washburn, M. P.; Van Nostrand, E. L.; Pratt, G. A.; Schwinn, M. K.; Daniels, D. L.; Yeo, G. W. SONAR Discovers RNA-Binding Proteins from Analysis of Large-Scale Protein-Protein Interactomes. *Molecular Cell* **2016**, *64* (2), 282–293. https://doi.org/10.1016/j.molcel.2016.09.003.
(19) Treiber, T.; Treiber, N.; Plessmann, U.; Harlander, S.; Daiß, J.-L.; Eichner, N.; Lehmann, G.; Schall, K.; Urlaub, H.; Meister, G. A Compendium of RNA-Binding Proteins That Regulate MicroRNA Biogenesis. *Molecular Cell* **2017**, *66* (2), 270-284.e13. https://doi.org/10.1016/j.molcel.2017.03.014.
(20) Caudron-Herger, M.; Rusin, S. F.; Adamo, M. E.; Seiler, J.; Schmid, V. K.; Barreau, E.; Kettenbach, A. N.; Diederichs, S. R-DeeP: Proteome-Wide and Quantitative Identification of RNA-Dependent Proteins by Density Gradient Ultracentrifugation. *Molecular Cell* **2019**, *75* (1), 184-199.e10. https://doi.org/10.1016/j.molcel.2019.04.018.
(21) Lambert, S. A.; Jolma, A.; Campitelli, L. F.; Das, P. K.; Yin, Y.; Albu, M.; Chen, X.; Taipale, J.; Hughes, T. R.; Weirauch, M. T. The Human Transcription Factors. *Cell* **2018**, *172* (4), 650–665. https://doi.org/10.1016/j.cell.2018.01.029.
(22) Edwards, A. L.; Garst, A. D.; Batey, R. T. Determining Structures of RNA Aptamers and Riboswitches by X-Ray Crystallography. *Methods Mol Biol* **2009**, *535*, 135–163. https://doi.org/10.1007/978-1-59745-557-2_9.
(23) Steiner, H. R.; Lammer, N. C.; Batey, R. T.; Wuttke, D. S. An Extended DNA Binding Domain of the Estrogen Receptor Alpha Directly Interacts with RNAs in Vitro. *Biochemistry* **2022**, *61* (22), 2490–2494. https://doi.org/10.1021/acs.biochem.2c00536.
(24) Hemphill, W. O.; Steiner, H. R.; Kominsky, J. R.; Wuttke, D. S.; Cech, T. R. Transcription Factors ERα and Sox2 Have Differing Multiphasic DNA and RNA Binding Mechanisms. *RNA* **2024**, rna.080027.124. https://doi.org/10.1261/rna.080027.124.
(25) Parsonnet, N. V.; Lammer, N. C.; Holmes, Z. E.; Batey, R. T.; Wuttke, D. S. The Glucocorticoid Receptor DNA-Binding Domain Recognizes RNA Hairpin Structures with High Affinity. *Nucleic Acids Research* **2019**, *47* (15), 8180–8192. https://doi.org/10.1093/nar/gkz486.
(26) Wieser, T. A.; Wuttke, D. S. Replication Protein A Utilizes Differential Engagement of Its DNA-Binding Domains to Bind Biologically Relevant ssDNAs in Diverse Binding Modes. *Biochemistry* **2022**, *61* (22), 2592–2606. https://doi.org/10.1021/acs.biochem.2c00504.
